# Brood-holding causes workers to pay attention to the queen in the carpenter ant *Camponotus japonicus*

**DOI:** 10.1101/383059

**Authors:** Kenji Hara

**Affiliations:** Life Science, Tokyo Gakugei University, 4-1-1, Nukui-Kitamachi, Koganei-shi, Tokyo 184-8501, Japan.

**Author notes:** **Name and address for correspondence:** Kenji Hara (Ph.D.), Life Science, Tokyo Gakugei University, 4-1-1, Nukui-Kitamachi, Koganei-shi, Tokyo 184-8501, Japan., e-mail.

**Keywords:** selective attention, olfaction, reiteration, social behavior, insect

## Abstract

**[Abstract]:** Brood accumulation, a fundamental behavior of offspring care in the carpenter ant *Camponotus japonicus*, is driven by alternation of ‘holding run’ and ‘empty-handed run’ behaviors. In the holding run, a worker holds a brood with her mandibles and carries it to the queen (holding run). After releasing it beside the queen, she hurries back to another brood (empty-handed run). To address the motivation for the brood-accumulation task, in this study, I observed these behaviors under experimental conditions. When workers performed the task in a situation that involved selection between their own and unfamiliar queens, they ran in significantly more restrictive ways during the holding run than during the empty-handed run. Hence, ‘holding’ represents a different motivational state than ‘empty-handed’. In a second experiment, the workers were suddenly presented with an unfamiliar floor during the task. Regardless of whether they were holding or empty-handed, their running traces on the familiar floor were simple, whereas on the unfamiliar floor they were more complex. These results show that holding workers would pay attention to the queen, exploiting cues on the floor to restrict their responses to the queen.

## [Introduction]

The behavioral complexity of insects has been demonstrated in behavioral and physiological studies of many species. Insects’ adaptive actions are based on reflexes and/or the internal patterns, modulated by their response to sensory stimuli. Motivation is a process that leads to the formation of behavioral intentions, i.e., it sets the aim of the behavior. In a behavior that is accomplished by combining multiple actions together, there is thought to be a different motivation for every action. For example, desert ants, *Cataglyphis fortis*, use path integration (PI) as their main mode of navigation (Wehner, 2003). Their foragers continuously measure directions and distances both when going out to a feeding site (outbound) and when returning to the nest (inbound), and then form outbound and inbound vectors, respectively, by integrating these two quantities. Utilization and switching of the vectors associated with the motivation allows the ants to perform the adaptive behavior (Collett *et al*., 1999; Merkle and Wehner, 2008). The appropriate change in motivation is necessary to achieve the goal resulting from a chain of actions.

In social insects, brood care by sterile workers plays important roles in maintenance of highly sophisticated communities. In the carpenter ant *Camponotus japonicus*, nestmates divide labor according to their ages, and brood care is the initial task after emergence. Nurse workers repeatedly carry scattered broods and accumulate them beside the queen (Hara, 2013), who affects brood growth by producing and secreting various substances (Holman, 2010; Motais de Narbonne *et al*., 2016). Therefore, brood-accumulation behavior is fundamental to the altruistic behavior responsible for rearing of kin broods by sterile workers.

The unit of brood-accumulation behavior consists of four sequential elements: (1) The worker recognizes a brood by touching it with the antennae, and then picks it up with her mandibles (‘pickup’). (2) The worker starts an inbound run while holding a brood (‘holding’). (3) The worker recognizes the queen by touching with the antennae, and then places the brood beside her (‘release’). (4) The worker turns her back on the queen and starts the outbound run (‘empty-handed’). Repetition of the unit results in gathering of broods near the queen. To address the physiological mechanisms underlying brood accumulation behavior, the workers performed tasks under two experimental conditions: a situation that involved selection between unfamiliar (UQ) and fostered queens (FQ) (Experiment 1), and a situation in which the familiar floor disappeared (Experiment 2). The data were analyzed to compare ‘holding’ and ‘empty-handed’ behaviors. I discuss the results from the standpoint of attention-like phenomena of workers engaging in the brood-accumulation task.

## [Materials and methods]

### Animals

Experiments were performed on the laboratory-reared workers and queens of the carpenter ant *Camponotus japonicus*. The laboratory colonies were found and maintained as previously described (Hara, 2002). Founding queens were collected from 2004 to 2016 in Tokyo, Japan. In order to obtain the healthy data, the workers and queens were used from the colonies within two years from the foundation.

To prepare the experimental workers, the pupae isolated from their birth colonies were removed from the cocoon and were incubated individually in the 96-well culture plate with U bottom (Ishii *et al*., 2005). Shortly after emergence, they were marked individually with cloth threads of different colors tied between their petiole and gaster and were transferred to the foster colonies (Hara, 2003). On acceptance by the natural ants and performance of the social activities, availability of those workers was decided for this experiment.

### Experimental design and trial procedure

*Experiment 1*: The brood-accumulation behaviors of the workers were recorded under the selective condition between the foster queen (FQ) and an unfamiliar queen (UQ). As details of the experimental procedure used for recording the behavior were given in a previous paper (Hara, 2003), only a brief outline is described below. An acrylic box was used for the test consisting of three rooms and the central space (Fig. 1A). Each room is connected to the central space by a doorway that allowed the workers free access to all rooms. A distance between the corners of the central space (red dots in Fig. 1A) was 2.7cm. Two queens and broods were put in the rooms of the box, Q1, Q2 and B, respectively. FQ was put in the room Q1 or Q2, and UQ was put in the other room. After acclimation for 15 min, the trial lasted 60 min. To avoid possible bias resulting from the UQ accidentally having the ‘colony labels’ similar to those of FQ, the preliminary check were performed every pair, with a control worker from the foster colony.

**Figure 1.**
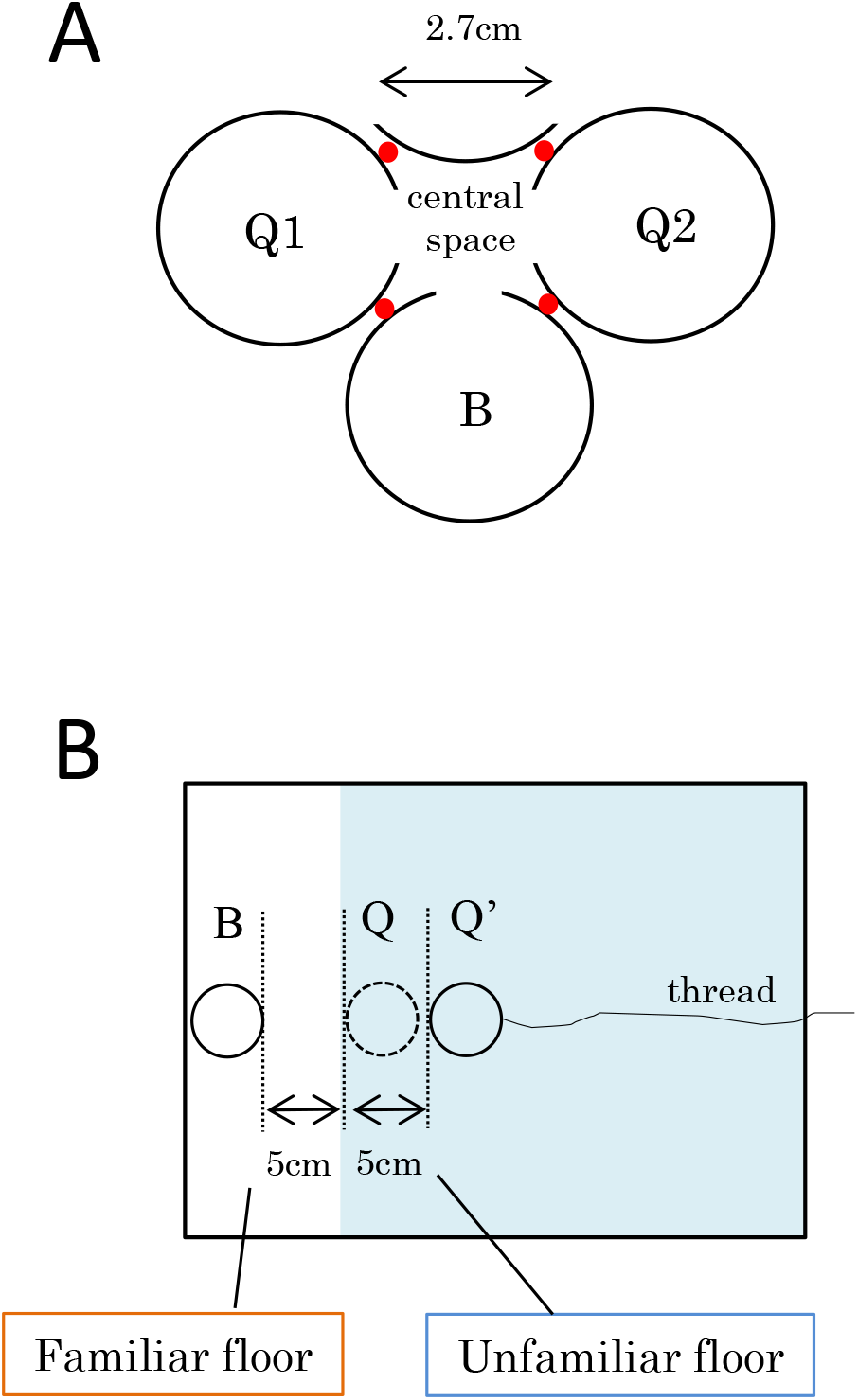
Schematic view of the experimental setups for (A) Experiment 1 and (B) Experiment 2. (A) A worker is allowed to carry a brood toward either the foster queen (in the one of Q1 or Q2) or an unfamiliar queen (in the other). The red dots indicate the datum-points for the array analysis. (B) A worker is allowed to carry broods in the dish “B” toward the queen in the dich “Q” in the preliminary test. In the middle of the final test, the Q-B distance is expanded by pulling the dish “Q” with a thread to the position of Q’. Subsequently, the worker has to run across the suddenly appeared (i.e., unfamiliar) floor to complete the task.

*Experiment 2*: To show an experimental worker the unfamiliar ground condition suddenly on the trial, the following apparatus was used (Fig.1B); Two plastic dishes (35 mm in diameter) were put in an acrylic box (252×345 mm). Broods were set in the one designated as “B” in Fig. 1B and queen was in the other “Q”. A thread was attached to the dish “Q” in order to operate it from outside the box. FQ was introduced in the apparatus and then a worker was allowed to explore the inside for 15 min. After finishing such acclimation, five broods from the foster colony were introduced into the dish “B” and then, the worker was put back into it. The behavior had been recorded for 60 min by the video camera since the worker picked a brood up.

The preliminary and the final tests were carried out sequentially with each worker. The distance between the dishes “Q” and “B” (B-Q distance) was constant at 5 cm through the preliminary trial. When the worker released the 3rd brood beside the queen in the final trial, the dish “Q” was moved away from the dish “B” to 10 cm (Fig. 1B). In this study, the zone to 5cm at linear distance from “B” is referred to as ‘familiar floor’ and ‘unfamiliar floor’ showed the zone more outer than it.

### Data analyses

All behaviors were recorded with the home video cameras (HDR-SR7, HDR-CX180, SONY; HDC-TM90, Panasonic; GZ-MG330, Victor). All data were captured at 30 frames per second. The video data were converted to AV1 format in the computers. To quantify movement, the position of the ant’s head was measured frame-by-frame using the motion analysis software, DIPP-MortionPro2D (Ditect, Japan). Two-dimensional coordinate data were saved as the excel file and were taken advantage of for subsequent analysis. All statistical calculations were performed with SPSS 22 and 23 (IBM, USA).

## [Results]

### Experiment 1

Effect of an unfamiliar queen as a distractor on brood-accumulation behavior

#### 1-1. Responses of workers to brood

In Experiment 1, I analyzed the responses of 149 laboratory-reared workers over 12 years, from 2004 to 2015. The results are summarized in Table 1A. Within 3 days after emergence, most workers were largely immobile (standstill). Six workers (21%) touched a brood with antennae, but five of them did not picked it up (i.e., they ignored it). At 4–7 days of age, there was an increase in the number of workers who held a brood: 13 (52%) held a brood and then released it beside another brood in the same room (gathering), but never carried it out of room “B”. The other five workers (20%) picked up a brood and left room “B” while holding it (transportation). After day 12, almost all workers performed transportation.

**Table 1.**
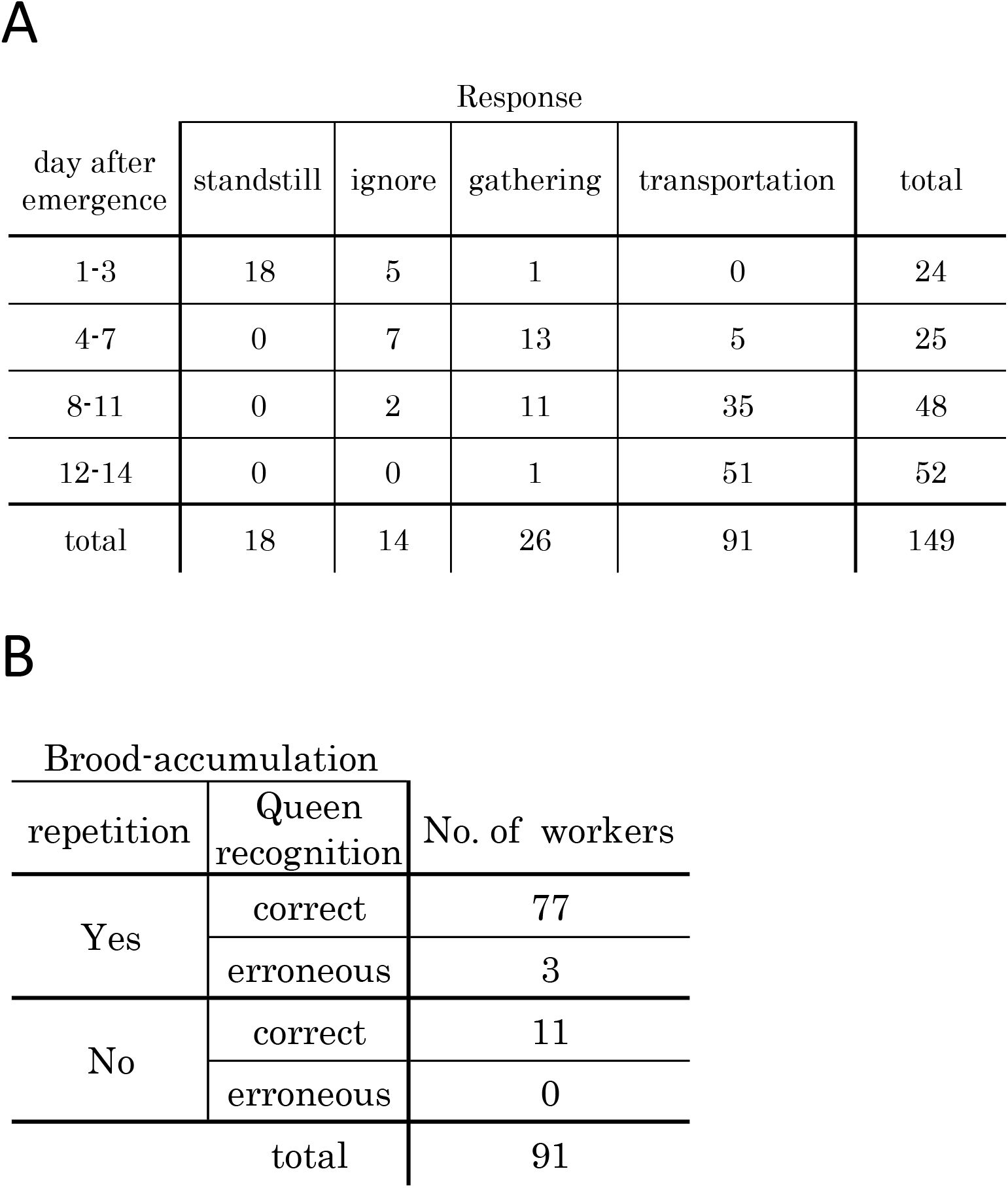
The workers tested in Experiment 1. (A) Their responses to brood. (B) Brood-accumulation behaviors of the workers identified as ‘transportation’.

According to both repetition of the activity and the ability to discriminate FQ from UQ, I performed further analysis of workers engaging in transportation; the results are summarized in Table 1B. Of 91 workers, 77 repeatedly carried their broods to the FQ. In this study, such brood-accumulation behavior characterized by repetition and FQ recognition is described as ‘regular’. Three workers brought some or all of broods to the UQ. Eleven workers delivered only one brood to the FQ and, after releasing it, did not leave the side of the FQ. No worker remained beside the UQ after incorrect delivery.

#### 1-2. Behaviors in the central space

For the 77 workers performing the regular task, I analyzed their trajectories in the central space for each holding and empty-handed run. A typical example is shown in Figure 2. During holding runs, the workers arrived at the FQ via restricted courses (Fig. 2A, C). By contrast, during empty-handed runs, the workers ran through a wider area and also seemed to be interested in the UQ (Fig. 2B, D). To quantify the special activities of workers, I defined an array by consolidating the position coordinates of the central space into a grid composed of 6 × 6 blocks; each block was 0.45 cm × 0.45 cm square (lower panels in Fig. 2C and 2D). Array analyses for 77 workers confirmed that the activity in the holding run was significantly more restricted, both spatially (P<0.01, Wilcoxon test; Fig. 3A) and temporally (P<0.01, test for the proportion; Fig. 3B) than in the empty-handed run.

**Figure 2.**
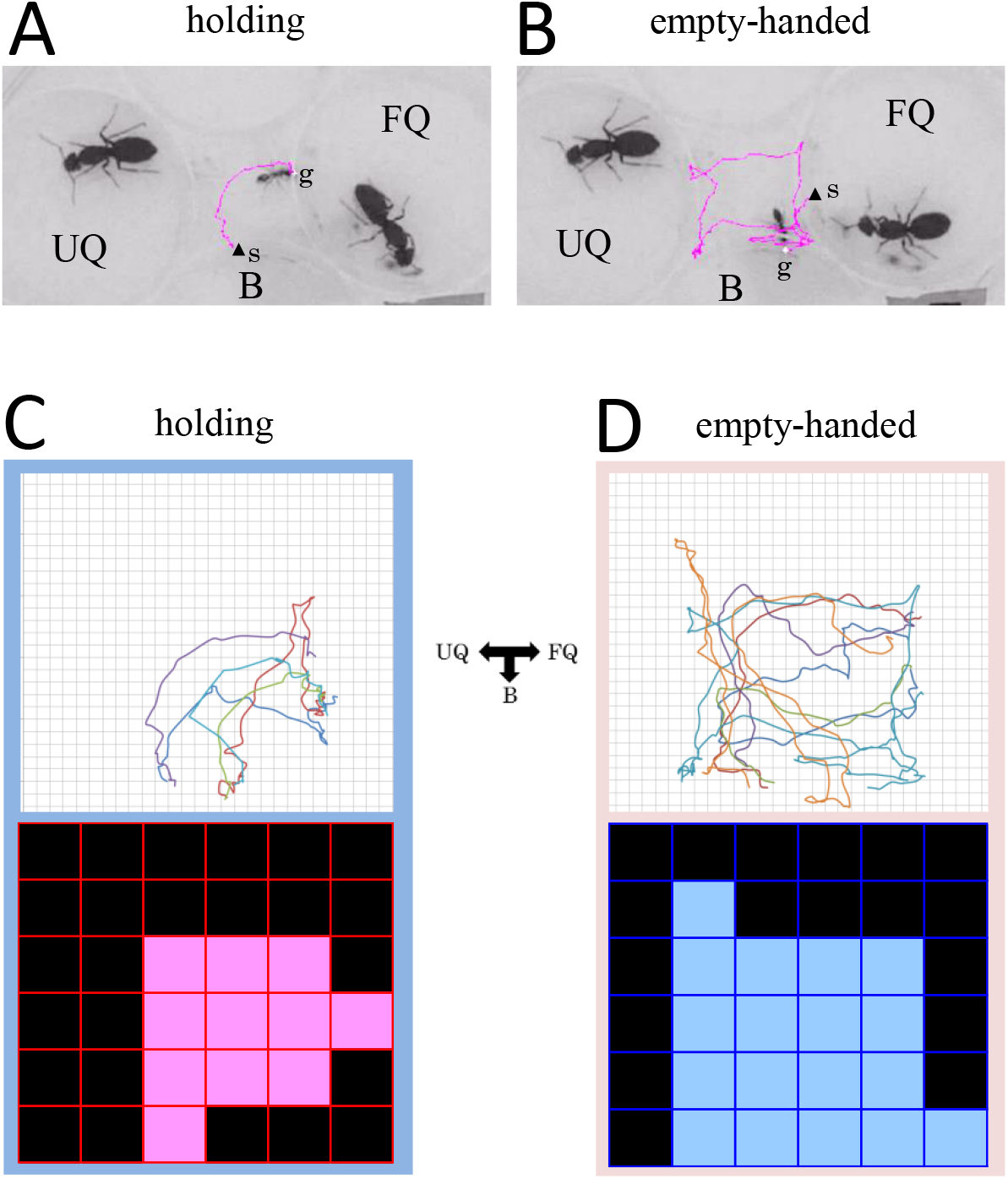
Behaviors in the central space in Experiment 1. Activities of a worker (A) during holding run for the 3^rd^ brood and (B) during subsequent empty-handed run. g=arrival point; s=starting point. For (C) the holding runs and (D) the empty-handed runs in one brood-accumulation task, tracks (upper panels) and the array analyses (lower panels) of a worker. The colors indicate different chains of running. FQ, the foster queen; UQ, an unfamiliar queen; B, the broods.

**Figure 3.**
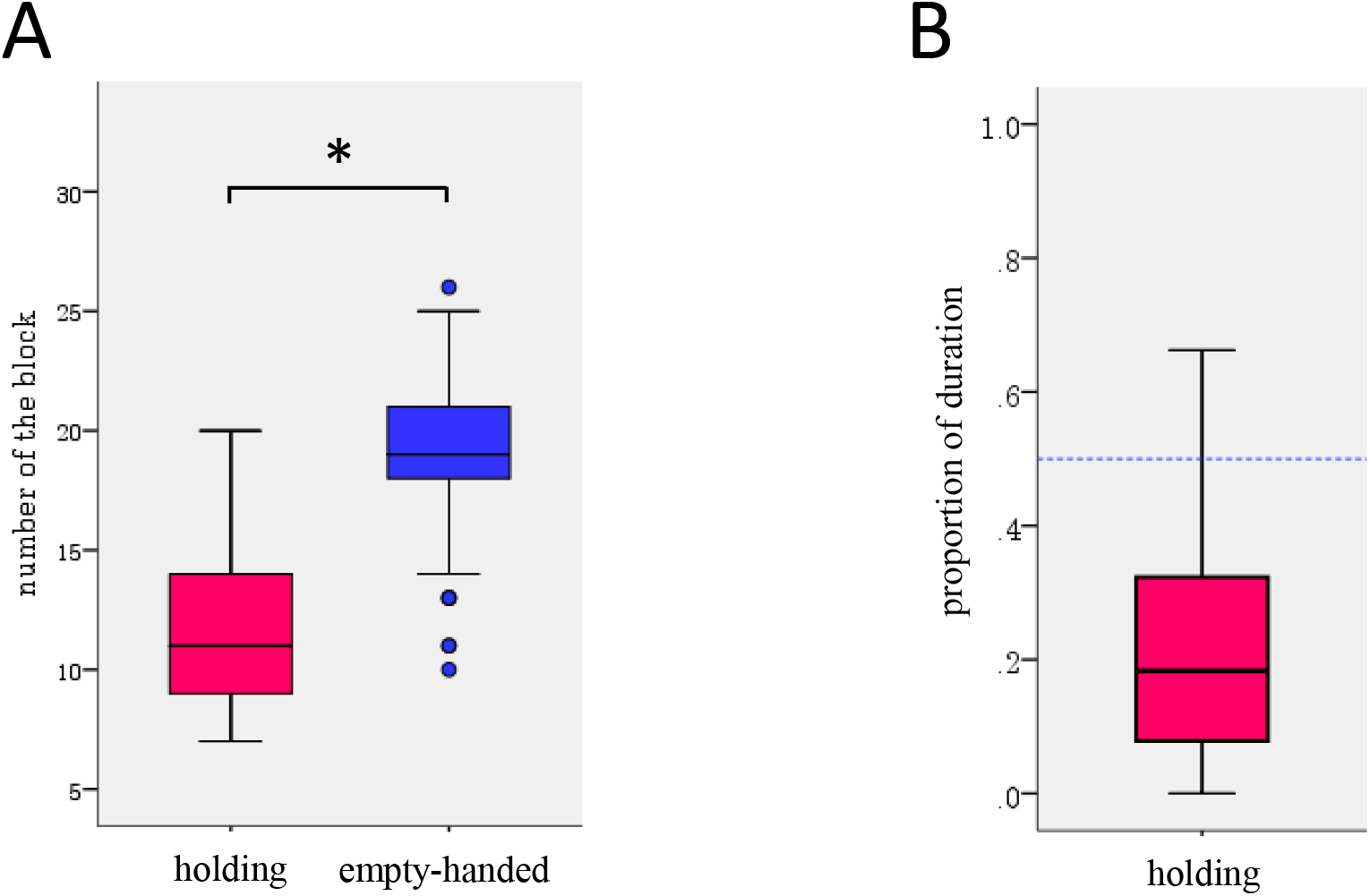
Results of array analyses from 77 workers in Experiment 1. (A) Box-plot representation of the spatial activities per task. Mean±s.d. = 11.91±3.56 in the holding runs, and 18.99±3.31 in the empty-handed runs. (B) The proportion of time for the holding runs in the central space per task. Mean±s.d. = 0.20±0.15. *P<0.01.

#### 1-3. Response of workers to queens

Queens induce behaviors in workers using pheromones. To determine whether the queen’s signals would influence repeatability in the brood-accumulation behavior, I analyzed the amounts of time each worker remained with the FQ. At the first delivery, the workers (n=77) clung to the FQ, and consequently stayed significantly longer in room “FQ” than during subsequent deliveries (P<0.01, Wilcoxon test; Fig. 4A). In particular, after the third visit, the workers left the room as soon as they released a brood beside the queen.

**Figure 4.**
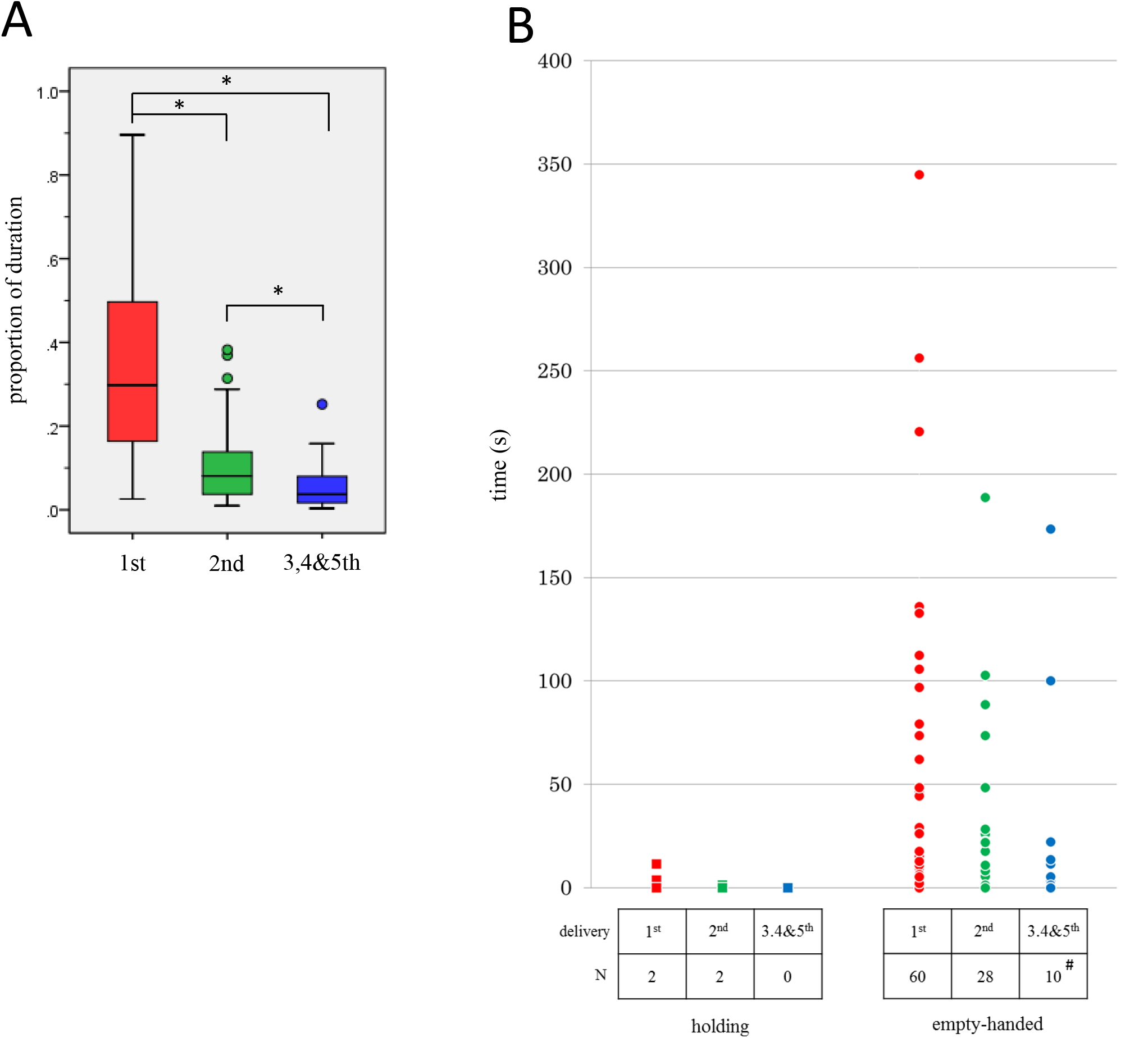
(A) The population of time for the workers to stay in the room “FQ” to the duration for the task. The data are presented as box plots every delivery. Because of shorter stays in each, the data at the 3^rd^, 4^th^ and 5^th^ deliveries are combined in one. Mean±s.d. = 0.33±0.21 at the 1^st^ delivery, 0.10±0.09 at the 2^nd^ and 0.05±0.05 at the 3, 4&5^th^. (B) Cumulative time per task to stay in the room “UQ”. N, the number of workers stepping in the room “UQ”; #, the number of workers stepping in the room “UQ” more than once at the 3^rd^, 4^th^ or 5^th^ deliveries. *P<0.01.

Few workers visited the UQ during the holding runs (Fig. 4B). Two workers at both the first and second delivery stepped into room “UQ” and immediately departed without touching the UQ. In the empty-handed runs, on the other hand, the frequency of UQ visits was much higher (Fig. 4B): 60 of 77 workers (78%) stepped into room “UQ” on the return trip after the first delivery. Some of them touched the UQ with their antennae a few times, and consequently spent longer in room “UQ”. At the 2^nd^ and subsequent deliveries, 28 (36%) and 10 (13%) of the empty-handed workers, respectively, visited the UQ (Fig. 4B).

### Experiment 2

Effect of an unfamiliar floor as a distractor on brood-accumulation behavior

#### 2-1. Responses of workers to brood

In Experiment 2, I analyzed responses to brood of 68 workers from laboratory-reared colonies over 6 years from 2011 to 2016. The results are summarized in Table 2A. In the preliminary test, 25 of 31 8–11-day-old workers and 36 of 37 12–14-day-old workers picked up a brood up and then carried it away from dish “B” (transportation). For workers engaging in transportation in the preliminary test, 21 of 25 8–11-day-old workers and all 36 12–14-day-old workers also transported a brood out of dish “B” in the final test. Ultimately, 57 workers (84%) had the ability to engage in brood transportation.

**Table 2.** The workers tested in Experiment 2. (A) Their responses to brood. ^(1)^, the number of workers analyzed in the preliminary test; ^(2)^, only the workers identified as ‘transportation’ in the preliminary test are tested in the final test. (B) Brood-accumulation behaviors of the workers engaging in ‘transformation’ in the final test.

| day after emergence | sample size <sup>(1)</sup> | Behavioral test | Response |  |  |  |
| --- | --- | --- | --- | --- | --- | --- |
|  |  |  | standstill | ignore | gathering | transportation |
| 8-11 | 31 | preliminary | 0 | 2 | 4 | 25 |
|  |  | final<sup>(2)</sup> | 0 | 1 | 3 | 21 |
| 12-14 | 37 | preliminary | 0 | 0 | 1 | 36 |
|  |  | final<sup>(2)</sup> | 0 | 0 | 0 | 36 |
| total | 68 | total<br>in the final tests |  |  |  | 57 |

Table 2
| Brood-accumulation |  |  |
| --- | --- | --- |
| repetition | mission | No.of workers |
| Yes | complete | 43 |
|  | incomplete | 6 |
| No | ----- | 8 |
| total |  | 57 |

I performed further analysis of those 57 workers; the results are summarized in Table 2B. 43 (75%) delivered five broods to the queen in both the preliminary and final tests. Six workers (11%) repeatedly carried broods to the FQ, but did not finish carrying all five broods in either or both tests (mission incomplete). Three of those workers strayed in the empty-handed run just after extending the B-Q distance and did not arrive at dish “B” before the time limit. Eight workers (14%) delivered only the first brood to the FQ and, after releasing it, did not leave the side of the FQ (no repetition).

#### 2-2. Behaviors on the unfamiliar floor

This experiment sought to determine whether workers engaging in the brood-accumulation task could continue their mission when they encountered a ground condition. The workers became familiar with the floor space between the broods and the queen by traversing it many times for brood delivery (Fig. 5A). In the holding run for the third brood in the final test, they ran straight to the queen (Fig. 5B). When the workers released the brood beside the queen, the B-Q distance was expanded to present an unfamiliar floor. In the subsequent empty-handed runs, the workers followed a meandering trace on the unfamiliar floor, whereas on the familiar floor they ran straight and easily arrived at dish “B” (Fig. 5C, upper panel in Fig. 5F). In the holding run for the fourth brood, the workers ran in an almost constant direction on the familiar floor but began to stray once they stepped onto the unfamiliar floor (Fig. 5D, upper panel in Fig. 5G). Their meandering runs were maintained until the delivery was completed, even if they ran back onto the familiar floor.

**Figure 5.**
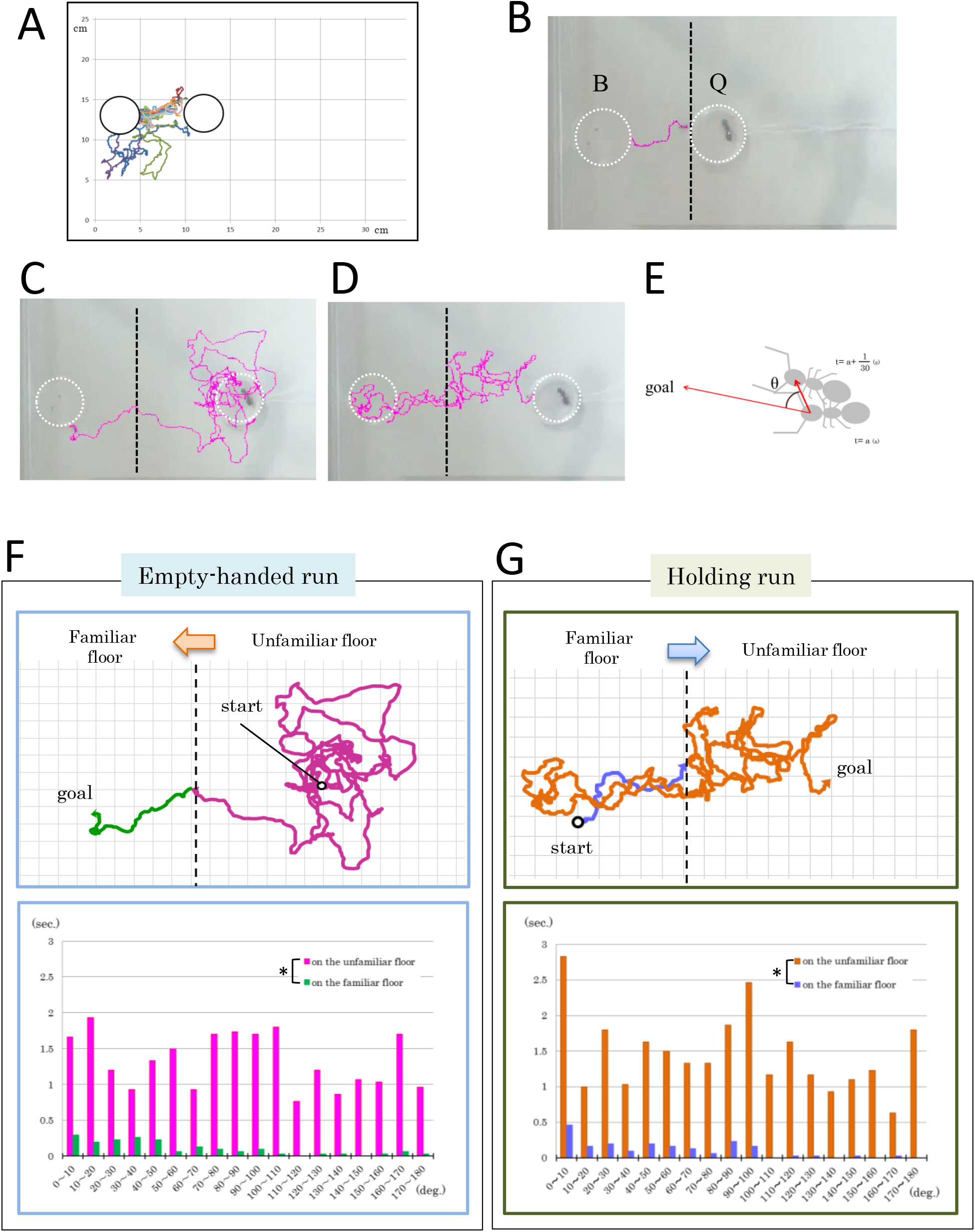
Behaviors of a worker in Experiment 2. (A) Tracks from beginning of the preliminary test to the ‘release’ of the 3^rd^ brood in the final test. The colors indicate different chains of running. (B) A track of the holding run for the 3^rd^ brood in the final test. B and Q denote the dishes (outlined) for the broods and the queen, respectively. A black short-dashed line indicates the boundary between the familiar and unfamiliar floors. (C) A track of the empty-handed run just after expanding the B-Q distance. (D) A track of the holding run for the 4^th^ brood in the final test. (E) Measurement of track angle (see text). Track-angle profiles in (F) the empty-handed run and (G) the holding run. Upper panels illustrate the trajectory of (C) or (D), respectively. The colors indicate the differences between the familiar and unfamiliar floor in each empty-handed or holding run. *P<0.01.

To quantify the effect of the ground condition on movement, I calculated frame-by-frame the track angle of the running trajectory, defined by the running vector and orientation to the goal; data were captured at 30 frames per second (Fig. 5E). The track angle histograms of empty-handed runs, composed of running profiles on the unfamiliar floor, showed that angles larger than 120 degrees were very frequent (lower panel in Fig. 5F). The profiles differed significantly between the unfamiliar and familiar floors (P<0.01, Wilcoxon test). In the holding runs, most of the track angles were smaller than 110 degrees until the workers stepped onto the unfamiliar floor. Once a worker crossed onto the unfamiliar floor, even if they ran back onto the familiar floor, angles larger than 120 degrees were very frequent (lower panel in Fig. 5G). Track angles differed significantly before and after stepping onto the unfamiliar floor (P<0.01, Wilcoxon test).

For all 43 workers that delivered all of five broods to the queen in both the preliminary and final tests (Table 2B), there were significant differences between the unfamiliar and familiar floor in both empty-handed and holding runs (P<0.01, Wilcoxon test; Fig. 6).

**Figure 6.**
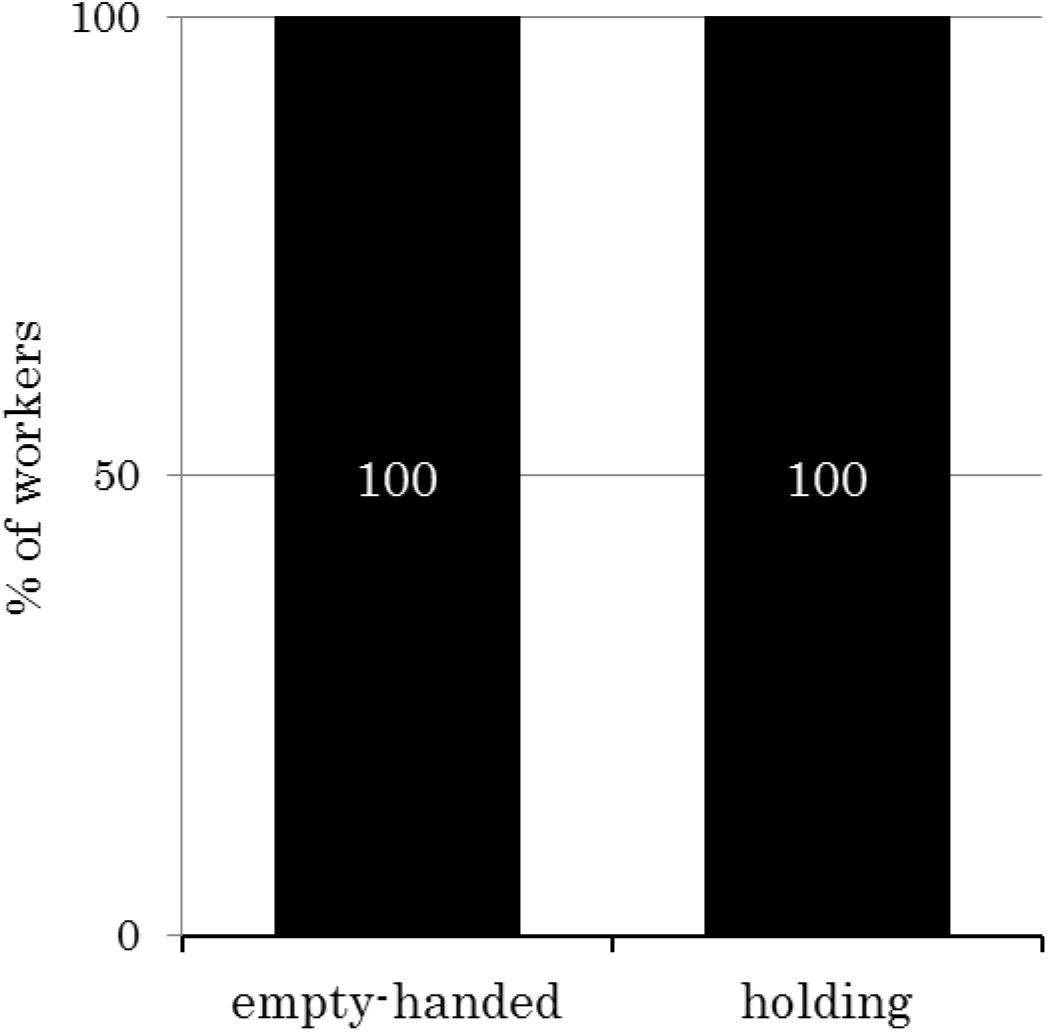
For each run, the percentage of workers whose angle profile is significantly different between the familiar and unfamiliar floor (P<0.01). N=43.

## [Discussion]

A *Camponotus* worker engaging in the brood-accumulation task alternates between two different states, ‘holding’ and ‘empty-handed’. In this study, I showed that the responsiveness of a worker to the queen differs significantly between these states, indicating that ‘holding’ represents a distinct physiological condition than ‘empty-handed’. The motivation for holding appears to be finding a queen for the brood being carried. This idea is supported by the observation that picking up a brood is a cue to start searching for the queen. On the other hand, the empty-handed state might also be associated with actual motivation. Immediately after releasing the brood beside the queen (however, that is not a case at the first brood), ‘empty-handed’ workers departed. Such quick conversion to the other state might be explained by a motivation to search for broods not already adjacent to the queen. Across insect species, internal representations of an animal’s own actions have been shown to constitute building blocks for appropriate behaviors (cf. Haberkern & Jayaramen, 2016). This idea supports the hypothesis that brood accumulation behavior might be driven by alteration of internal representations associated with the holding and empty-handed states.

For workers engaging in the brood-accumulation task, reaching the queen is essential. To address this issue, I presented workers engaging in the task with an unfamiliar floor by increasing the distance between the broods and the queen. Regardless of whether they were holding or empty-handed, the workers strayed on the unfamiliar floor. If workers recognize the distance from ‘start’ to ‘goal’ and exploit it as a clue to achieve their tasks, then they should stray on the latter half of their trip. Consequently, information about running distance would not be available for brood-accumulation behavior.

Following chemical signposts is the most straightforward method for reaching the destination. Pheromone trails are well-studied systems used to guide foragers between the nest and food sources. Previous research on pharaoh’s ants (*Monomorium pharaonis*) demonstrated that pheromone trails include polarity information (Jackson *et al*., 2004). Inside the dark nest, multiple chemicals including trails are attached to the tunnel surface because workers walk back and forth within the narrow space. In addition, the queen moves and does not stay in the same place. Under such circumstances, it would seem impossible for workers to distinguish the polarity information included in the trails and exploit it to guide their round-trip behavior. In this study, no workers quit their task because of the unfamiliar floor (i.e., the presumed disappearance of trail pheromone). Thus, it would be difficult to explain brood-accumulation behavior in terms of route-following guided by pheromonal signals.

Marking is also well accepted as a type of chemical sign to operate in the pinpoint. Home-range marking, for example, plays a crucial role in the vicinity of the nest to define the property of the colony, and both indicate the nest entrance to returning workers and provides defense against non-nestmates (Hölldobler and Wilson, 1990). Such markings often consist of non-volatile compounds from the ant’s cuticle surface, and therefore must be perceived by direct antennal contact (Lenoir *et al*., 2009). This evidence supports the hypothesis that in order to achieve the brood-accumulation task, workers must enhance orientation performance with the help of some marking material that acts like a footprint. This idea is consistent with this study’s finding that workers strayed, but never gave up on accomplishing the task on the unfamiliar floor.

Recognition of and attraction to the queen by workers is generally mediated by chemoreception (Hölldobler and Wilson, 1990). Therefore, the ability to select the appropriate information from a complex mixture of chemical components is important for the brood-accumulation task, as well as other social behaviors of *Camponotus* workers. Attention is a physiological mechanism by which animals select useful information by filtering the stimuli from the environment (Aston-Jones *et al*., 1999). Human studies of attention have shown that this selection is controlled either in a voluntary, top-down way or in a passive, bottom-up way (Motter, 1994; Theeuwes, 2010). Recent studies in several insects have revealed attention-like processes involved in selection of one among many visual and auditory stimuli (van Swindern and Greenspan, 2003; Spaethe *et al*., 2006; Wiederman and O’ Carroll, 2013; de Bivort and van Swindern, 2016), and careful comparisons of the mechanistic similarities between insects and primates have been carried out (cf. Nityananda, 2016). Very little work, however, has been published on olfactory attention. In this study, I showed that brood-holding can lead a worker to pay attention to the queen’s label. It remains to be demonstrated whether this process is elicited by attentional capture or an endogenous cueing procedure. Mature workers that are isolated from the queen immediately after their emergence do not perform the brood-accumulation behavior (Hara, 2003), indicating that ‘social association’ might be necessary to induce some neuronal modulations associated with the motivation for and/or the alteration of the holding and empty-handed states. To address the mechanism of olfactory attention, it might be useful to exogenously induce brood-accumulation behavior in such isolated workers.

## [Acknowledgments]

I wish to express my deepest thanks to three undergraduate members in my laboratory, Chie Sato (2005–2007), Sho Suzuki (2011–2013) and Ayane Tanaka (2012–2014), for invaluable assistance with the collection of data. In fifteen years from 2003 to 2017, a total of fifteen undergraduate students belonging to my laboratory helped me to take care of the ant colonies. I also thank them. This study was supported financially by Tokyo Gakugei University.

